# Mutant KRAS modulates colorectal cancer cells invasive response to fibroblast-secreted factors through the HGF/C-MET axis

**DOI:** 10.1101/2021.11.16.468815

**Authors:** Patrícia Dias Carvalho, Flávia Martins, Susana Mendonça, Andreia Ribeiro, Ana Luísa Machado, Joana Carvalho, Maria José Oliveira, Sérgia Velho

## Abstract

Genetic alterations influence the malignant potential of cancer cells, and so does the tumor microenvironment. Herein, we combined the study of KRAS oncogenic effects in colorectal cancer cells with the influence of fibroblasts-derived factors. Results revealed that mutant KRAS regulates cell fate through both autonomous and non-autonomous signaling mechanisms. Specifically, processes such as proliferation and cell-cell aggregation were autonomously controlled by mutant KRAS independently of the stimulation with fibroblasts conditioned media. However, cancer cell invasion revealed to be a KRAS-dependent non-autonomous effect, resulting from the cooperation between fibroblasts-derived HGF and mutant KRAS regulation of C-MET expression. C-MET downregulation upon KRAS silencing rendered cells less responsive to HGF and thus less invasive. Yet, in one cell line, KRAS inhibition triggered invasion upon stimulation with fibroblasts conditioned media. Inhibition of PIK3CA oncogene did not promoted invasion, thus showing a KRAS-specific effect. Moreover, the invasive capacity also depended on the HGF-C-MET axis. Overall, our study awards oncogenic KRAS an important role in modulating the response to fibroblast-secreted factors either by promoting or impairing invasion, and depicts the HGF-C-MET axis as a putative therapeutic target to impair the invasive properties of mutant KRAS cancer cells.

**Significance:** Targeting mutant KRAS cancers is an urgent clinical need. HGF-C-MET axis inhibition arises as a possible strategy to target mutant KRAS CRC, both primary and metastatic tumors.

**Additional information:** *Financial support:* This work was supported through FEDER funds through the Operational Programme for Competitiveness Factors (COMPETE 2020), Programa Operacional de Competitividade e Internacionalização (POCI), Programa Operacional Regional do Norte (Norte 2020), European Regional Development Fund (ERDF), and by National Funds through the Portuguese Foundation for Science and Technology (FCT) (PTDC/MED-ONC/31354/2017). PDC is a PhD student from Doctoral Program in Pathology and Molecular Genetics from the Institute of Biomedical Sciences Abel Salazar (ICBAS, University of Porto) and she is funded through a PhD fellowship (SFRH/BD/131156/2017) awarded by the FCT. FM is a PhD student from Doctoral Program in Biomedicine from the Faculty of Medicine of the University of Porto and she is funded through a PhD fellowship (SFRH/BD/143669/2019) awarded by the FCT. SM is a PhD student from Doctoral Program in Biomedicine from the Faculty of Medicine of the University of Porto and she is funded through a PhD fellowship (SFRH/BD/143642/2019) awarded by the FCT. AR is a junior researcher hired by IPATIMUP under the CaTCh project funded by FEDER and FCT (POCI-01-0145-FEDER-031354). ALM is a PhD student from Doctoral Program in Biomedicine from the Faculty of Medicine of the University of Porto and she is funded through a PhD fellowship (2020.08932.BD) awarded by the FCT. MJO is principal researcher at INEB. SV is hired by IPATIMUP under norma transitória do DL n.º 57/2016 alterada pela lei n.º 57/2017.

## Introduction

Accounting for more than 1.9 million cases and 935,000 deaths in 2020, colorectal cancer (CRC) ranks third in terms of incidence and steps up to the second place in terms of mortality (1). The estimated 5-year overall survival (OS) rate for all disease stages is 65%. However, approximately 50% of the patients progress to metastatic disease, and their 5-year OS is just of 14% (2). Importantly, the frequency of KRAS mutations within the metastatic subset can reach 50% of the cases (3), associating with poor prognosis. Accordingly, CRC patients with KRAS-mutant tumors have worse overall survival, increased incidence of lung, bone and brain metastases (4–6) as well as increased recurrence and shorter post-recurrence survival after hepatectomy for colorectal liver metastasis (7). Moreover, mutant KRAS CRCs are also more refractory to standard treatments and non-responsive to anti-EGFR therapies (8–10). Adding another line of importance, the occurrence of KRAS *de novo* mutations have been shown to be major events in anti-EGFR acquired resistance (11). Therefore, finding ways to improve the prognosis of mutant KRAS CRC patients is an urgent clinical need. As such, great efforts have been made along the years to specifically target KRAS or its downstream signaling effectors. Despite the renewed hope brought by the recent approval of the specific KRAS G12C inhibitor, this mutation occurs in only 3% of CRC cases (12). Therefore, KRAS mutant CRC patients, still lack targeted and efficient therapeutic options.

The understanding of cancer as a complex ecosystem and the recognition of the importance of the tumor microenvironment (TME) in disease progression and therapy response, changed the view of cancer therapy. Many therapies targeting the TME have been/are being developed and represent promising approaches for cancer treatment (13). Hence, the successful definition of therapeutic targets relies on the study of the communication of cancer cells with the surrounding TME components. Currently, it has been acknowledged that KRAS oncogenic effects are highly related to its capacity to modulate the TME (14–18). Within the TME, cancer- associated fibroblasts (CAFs) represent a heterogeneous population that can have diverse origins, with the majority originating from local normal fibroblasts that became “activated” (CAFs) through a variety of mechanisms. For instance, tumor cells themselves and other cells residing in the TME may secrete soluble factors, for example transforming growth factor-β (TGF-β), platelet-derived growth factor (PDGF) and interleukin (IL)-6, which can activate fibroblasts, generating CAFs. These are characterized by the expression of markers such as α-smooth muscle actin (α-SMA), fibroblast activation protein-α (FAP-α) or fibroblast-specific protein-1 (FSP-1/S100A4). In their turn, CAFs secretome is rich in growth factors, cytokines, exosomes, and metabolites that are essential to support cancer cells and other components of the TME, dictating tumor progression (19,20). In CRC, CAFs represent the main stromal constituents (19) and have long been recognized as essential modulators of cancer cells invasion and metastasis (21). Importantly, CAFs markers are found at the invasive front of CRC cases (22).

Despite the accumulating data elucidating the communication of KRAS mutant cancer models with the TME components, works approaching the interplay between KRAS mutant cancer cells and fibroblasts are still scarce, particularly in CRC. Herein, we questioned: is mutant KRAS sufficient to promote the malignant features of CRC or does it cooperate with fibroblasts-derived factors to induce them? Are fibroblasts able to support CRC malignant features in the context of KRAS inhibition, thus functioning as possible mechanism of resistance to KRAS-targeting? Our results revealed that mutant KRAS regulates proliferation and cell-cell aggregation through an autonomous signaling mechanism. However, invasion was shown to result from the cooperation between KRAS oncogenic activity and fibroblasts-secreted factors. Importantly, we showed that KRAS silencing prompted opposite effects, decreasing or increasing the invasive capacity in response to fibroblast-secreted factors, depending on the cell line. Mechanistically, we disclosed the KRAS-C-MET-HGF axis underlying the invasive responses.

## Material and methods

### Cell culture

CRC cell lines HCT116, HCT15 and SW480 as well as CCD-18Co normal colon fibroblasts were purchased from American Type Culture Collection (USA). SW620 and LS174T were kindly provided by Dr. Ragnhild A. Lothe (Oslo University Hospital).

HCT116, SW480 and HCT15 were cultured in RPMI 1640 medium (Gibco, Thermo Fisher Scientific) and SW620, LS174T and CCD-18Co were cultured in DMEM medium (Gibco, Thermo Fisher Scientific). For all cell lines the respective medium was supplemented with 10% fetal bovine serum (Hyclone) and 1% penicillin–streptomycin (Gibco, Thermo Fisher Scientific). Cells were maintained at 37°C in a humidified atmosphere with 5% CO_2_.

### CCD-18Co conditioned media production

For conditioned media production, the same number of cells was plated in two T75 culture flasks and cultured until approximately 90% of confluence. When the desired confluence was reached, cells were washed twice with PBS 1x and new media was added. In one of the flasks, corresponding to “normal-like” fibroblasts, DMEM supplemented only with 1% penicillin/streptomycin was used. In the other flask, corresponding to “activated-fibroblasts” the same volume of DMEM supplemented with 1% penicillin/streptomycin and 10 ng/mL rhTGFβ1 (ImmunoTools GmbH) was added. In parallel the same conditions-DMEM+1% penicillin/streptomycin and DMEM+1% penicillin/streptomycin+ 10 ng/mL rhTGFβ1-were employed using culture flasks with no cells, serving as controls. After four days in optimal culture conditions, conditioned media (CM) were collected, centrifuged, filtered through a 0.2 μm filter and stored at −20ºC (for a maximum of 6 months). Cells were tripsinized and counted to assure an equivalent number of cells in both conditions. Total protein was extracted, and fibroblasts’ activation was confirmed through the evaluation of α-SMA expression by western blot.

### Protein depletion from CCD-18Co conditioned media

Protein depletion from the CM was performed according to the protocol described by Gong et al., 2020 (23). Briefly, CM was frozen at −80°C and thawed at 37°C in three consecutive cycles of 15 minutes plus 15 minutes. Then, CM was passed through a 0.2 μm filter and centrifuged, according to manufacture instructions, in Amicon Ultra-15 centrifugal filter units with a 3-kDa cutoff, to allow protein removal. To evaluate proper protein depletion an additional step was performed. Total protein was quantified using the DCProtein assay kit (Bio-Rad) and 20 μg were resolved on sodium dodecyl sulphate-polyacrylamide gel electrophoresis (SDS-PAGE) under denaturing conditions and transferred to Hybond ECL membrane (Amersham Biosciences, GE Healthcare). Membrane was stained with Ponceau S solution (Sigma-Aldrich).

### siRNA transfection

CRC cell lines were seeded in six-well plates (see Supplementary Table S1 for the number of plated cells) and transfected after approximately 16 hours, using Lipofectamine RNAiMAX (Invitrogen, Thermo Fisher Scientific) in reduced-serum Opti-MEM medium (Gibco, Thermo Fisher Scientific). Gene silencing was achieved with ON-TARGETplus SMARTpool small interfering RNA (siRNA) specific for KRAS (L-005069-00-0010), C-MET (L-003156-00-0005) and HER3 (L-003127-00-0005) at a final concentration of 10 nM, and PIK3CA (L-003018-00-0010) at a final concentration of 50 nM, all from Dharmacon. As a negative control (siCTRL), a non-targeting siRNA (D-001810-01-50; ON-TARGETplus Non-targeting siRNA #1) was used. Subsequent assays were performed upon 72 hours of cell transfection for KRAS, C-MET and PIK3CA and 48 hours for HER3. Silencing efficiency was monitored by western blot.

### Cell cycle evaluation

Cell cycle was assessed using Click-iT™ Plus EdU Alexa Fluor™ 647 Flow Cytometry Assay kit (Invitrogen, ThermoFisher Scientific). Briefly, after 72 hours of transfection, siCTRL and siKRAS cells were washed and treated with serum-free CM (DMEM, DMEM+rhTGFβ1, Fibroblasts and Fibroblasts+rhTGFβ1). After 24 hours, EdU (5-ethynyl-2′-deoxyuridine) was added to each condition at a final concentration of 10 μM and incubated during 1 hour. Cells were trypsinized and approximately 2×10^5^ cells were fixed using the Click-iT™ fixative for 15 minutes at room temperature (RT), protected from light. After one wash in 1% bovine serum albumin (BSA; NZYtech) in PBS 1x, cells were resuspended in 1x Click-iT™ saponin-based permeabilization and wash reagent (washing buffer) and incubated for 10 minutes at RT, in the dark. The Click-iT™ Plus reaction cocktail was prepared according to the proportions indicated by the manufacturer to a final volume of 200 μL per sample, and incubated during 30 minutes at RT, in the dark. Finally, cells were washed once with Click-iT™ washing buffer and once in PBS 1x. Pelleted cells were resuspended in 100 μL of RNase A (Life Technologies, ThermoFisher Scientific) prepared in PBS 1x (final concentration of 100 μg/mL) and incubated for 15 minutes at 37ºC. Following the last incubation, 100 μL of 1% BSA were added to each sample. Before acquisition in a FACSCanto II cytometer (BD Biosciences), 0.5 μg/mL of propidium iodide (Sigma-Aldrich) was added to each sample. Results were analyzed in FlowJo software version 10.

### Slow aggregation assay

After 72 hours of transfection, siCTRL and siKRAS cells were trypsinized. A cell suspension of 1×10^5^ cells/mL in CM (DMEM, DMEM+rhTGFβ1, Fibroblasts and Fibroblasts+rhTGFβ1) supplemented with 10% FBS (Hyclone) was prepared. From this cell suspension, 200 μL/well were plated in triplicate into a 96-well plate pre-coated with 50 μL of bacto-agar (BD Biosciences) in PBS1x (0,66% w/v). The formation of aggregates was evaluated at 24 and 48 hours under a Leica DMi1 inverted microscope using the 5x objective.

### Enzyme-linked immunosorbent assay (ELISA)

The levels of known pro-invasive factors present in the CM from non-activated and activated fibroblasts, were quantified using ELISA kits following the manufacturer instructions. Specifically, HGF was quantified using a kit from RayBiotech and total TGFβ1 using a kit from Legend Max (BioLegend).

### Western blotting

Total protein was extracted using ice cold RIPA lysis buffer (50 mM Tris HCl; 150 mM NaCl; 2 mM EDTA; 1% IGEPAL CA-630; pH=7.5) supplemented with a protease inhibitor cocktail (Roche) and a phosphatase inhibitor cocktail (Sigma-Aldrich). Protein concentration was determined using the DCProtein assay kit (Bio-Rad) and analyzed by Western blotting. Briefly, protein extracts were resolved on SDS-PAGE gels under denaturing conditions and transferred to Hybond ECL membranes (Amersham Biosciences, GE Healthcare). Membranes were blocked for 1 hour at RT and incubated overnight with the respective primary antibody (Supplementary Table S2) at 4ºC with agitation. After 1 hour incubation at RT with the respective anti-mouse (NA931, GE Healthcare) or anti-rabbit (#7074, Cell Signaling) HRP-conjugated secondary antibodies, bands were detected using ECL (Bio-Rad) and autoradiography film (Amersham Biosciences, GE Healthcare) exposure. Band intensity was quantified using ImageJ.

### Phospho-RTK array

After 72 hours of transfection with KRAS siRNA, the medium was removed, cells were washed and treated with serum-free control and activated-fibroblasts CM (DMEM+rhTGFβ1 and Fibroblasts+rhTGFβ1) for 24 hours. Total protein was extracted using the Human Phospho RTK-array kit (R&D Systems) lysis buffer, supplemented with protease inhibitor cocktail (Roche), according to manufacturer instructions. Protein concentration was determined using the DCProtein assay kit (Bio-Rad) and 300 μg of total protein were incubated overnight at 4ºC with agitation with pre-blocked array membranes. After washing, array membranes were incubated with Anti-Phospho-Tyrosine-HRP Detection Antibody (provided in the kit) for 2 hours at RT with agitation. Following another round of washes, spots were revealed with the kit Chemi Reagent Mix and autoradiography film (Amersham Biosciences, GE Healthcare) exposure.

### Matrigel invasion assay

Invasion assays were performed using 24-well Matrigel invasion chambers (Corning BioCoat Matrigel Invasion Chamber, 8.0 μm PET Membrane). Briefly, cells were detached with trypsin and 1×10^5^ cells resuspended in complete culture medium were seeded in the upper compartment of pre-hydrated inserts. In the bottom compartment, CM supplemented with 10% FBS was used as chemoattractant. The plate was incubated in optimal culture conditions for 24 hours (in the case of HCT116, HCT15 and LS174T) or 48 hours (in the case of SW480 and SW620). Non-invasive cells were removed from the upper part of the filter with a cotton swab and filters were washed in PBS 1x. Invasive cells were fixed in ice-cold methanol for 20 minutes and mounted in slides using Vectashield with DAPI (Vector Laboratories) mounting medium. Invasion was quantified by counting the total number of invasive nuclei under a Leica DM2000 fluorescence microscope.

### HGF supplementation and neutralization

DMEM+rhTGFβ1 control medium was supplemented with 50 ng/mL rhHGF (ImmunoTools GmbH). To achieve HGF neutralization, Fibroblasts+rhTGFβ1 medium was incubated with 1 μg/mL of anti-human HGF antibody or with the respective mouse IgG1 isotype control (both from R&D Systems) for 1 hour at 37ºC.

### Statistical analysis

Statistical analyses were performed using the GraphPad Prism Software (version 7.00). Results are expressed as mean ± standard deviation (SD) and the specific performed test, as well as the number of independent biological replicates, are referred on each figure legend.

## Results

### Mutant KRAS exerts both autonomous and non-autonomous oncogenic effects

To determine whether fibroblasts secreted factors potentiate the oncogenic effects of mutant KRAS or provide pro-tumor signals allowing cells to tolerate KRAS inhibition, we performed a series of functional studies such as cell cycle assessment, cell-cell aggregation, and cell invasion. To do so, we used a panel of five mutant KRAS CRC cell lines (Supplementary Table S3) in which KRAS expression was silenced through siRNA (see Supplementary Fig.S1).

To evaluate if fibroblast-secreted factors were sufficient to sustain mutant KRAS cell proliferation, control (siCTRL) and KRAS silenced (siKRAS) cells were challenged with serum-free control CM (DMEM and DMEM+rhTGFβ1) as well as with CM from non-activated/normal-like and activated/CAF-like fibroblasts (Fibro. and Fibro.+rhTGFβ1, respectively; see Supplementary Fig. S2 for confirmation of activation). Notably, all siKRAS cell lines displayed a reduction in the number of cells when compared to siCTRL cells. Treatment with CM from non-activated or activated fibroblasts (Fibro. and Fibro.+rhTGFβ1, respectively) neither affected the number of cells in the siCTRL condition nor it rescued the reduction observed upon KRAS silencing (Fig. 1A). Cell cycle analysis revealed that in HCT15, SW620 and SW480 cells treated with control CM (DMEM and DMEM+rhTGFβ1), siKRAS promoted a significant increase in the percentage of cells arrested in G1 (Fig. 1B). Only in HCT15, exposure of siKRAS cells to factors secreted by activated fibroblasts (Fibro+rhTGFβ1 CM) led to the reduction of the percentage of cells in G1 (Fig. 1B), without affecting the total number of cells (Fig. 1A). No cell cycle alterations were found in HCT116 and LS174T cells upon KRAS silencing or stimulation with fibroblasts CM. In addition, treatment with fibroblasts CM did not induce any cell cycle alterations in siCTRL cells in neither of the cell lines (Fig. 1B).

**Figure 1.**
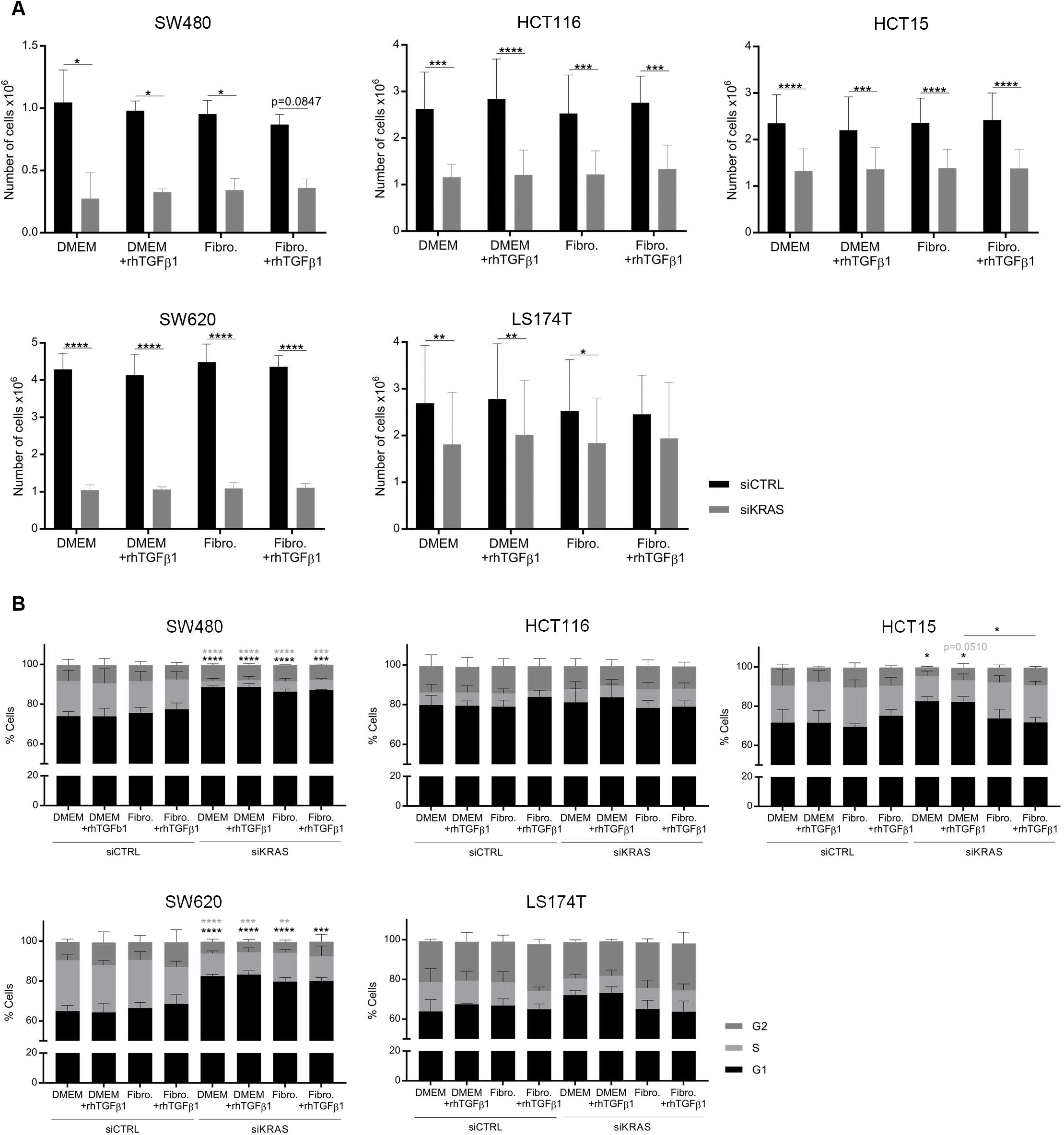
Fibroblast CM is not enough to rescue the effects of KRAS silencing on cell number and proliferation. Serum-free CM from fibroblasts and rhTGF-β-activated fibroblasts and the respective controls (DMEM and DMEM+rhTGFβ1) were used to treat siCTRL and siKRAS cells for 24 hours. **A**, KRAS silencing results in a decrease in the number of cells that is not affected by the treatment with fibroblasts-CM. **B**, HCT116 and LS174T cell lines shown no differences in cell cycle between conditions. HCT15 siKRAS treated with DMEM and DMEM+rhTGFβ1 show a significative increase in the percentage of cells in G1 phase when compared to the respective siCTRL cells. When comparing this condition with the Fibro.+rhTGFβ1, there is a decrease in the percentage of cells in G1 phase. Both in SW620 and SW480 all conditions of siKRAS cells show an increase in the percentage of cells in G1 phase acompained with a significative decrease in S phase (with the exception of SW620 siKRAS treated with Fibro.+rhTGFβ1 CM). Values are ploted as mean ±SD of at least 3 biological replicates and statistical significance was evaluated using the two-way ANOVA considering repeated measures by both factors, with Tukey’s multi comparison test (*p≤0.05; **p≤0.01; ***p≤0.001; ****p≤0.0001).

To infer the impact of fibroblasts secreted factors on mutant KRAS cancer intercellular adhesion, a classical slow aggregation assay was performed. The data showed that the panel of cell lines have heterogenous cell-cell aggregation capacities, that in all cases are independent of fibroblasts CM treatment (Fig. 2 and Supplementary Fig. S3). In particular, SW480 siCTRL cells did not form aggregates and neither siKRAS nor the fibroblasts CM reverted this state. KRAS silencing in HCT116 and HCT15 cells resulted in an increase of the aggregation capacity but no effect was observed upon culture with CM from non-activated or activated fibroblasts. SW620 siCTRL formed very loose aggregates, but upon KRAS silencing, and independently of treatment with CM of fibroblasts, cells formed smaller but more compact aggregates when compared with siCTRL. Despite forming aggregates, LS174T cells aggregation capacity was not affected neither by KRAS silencing nor by the fibroblasts CM (Fig. 2 and Supplementary Fig. S2).

**Figure 2.**
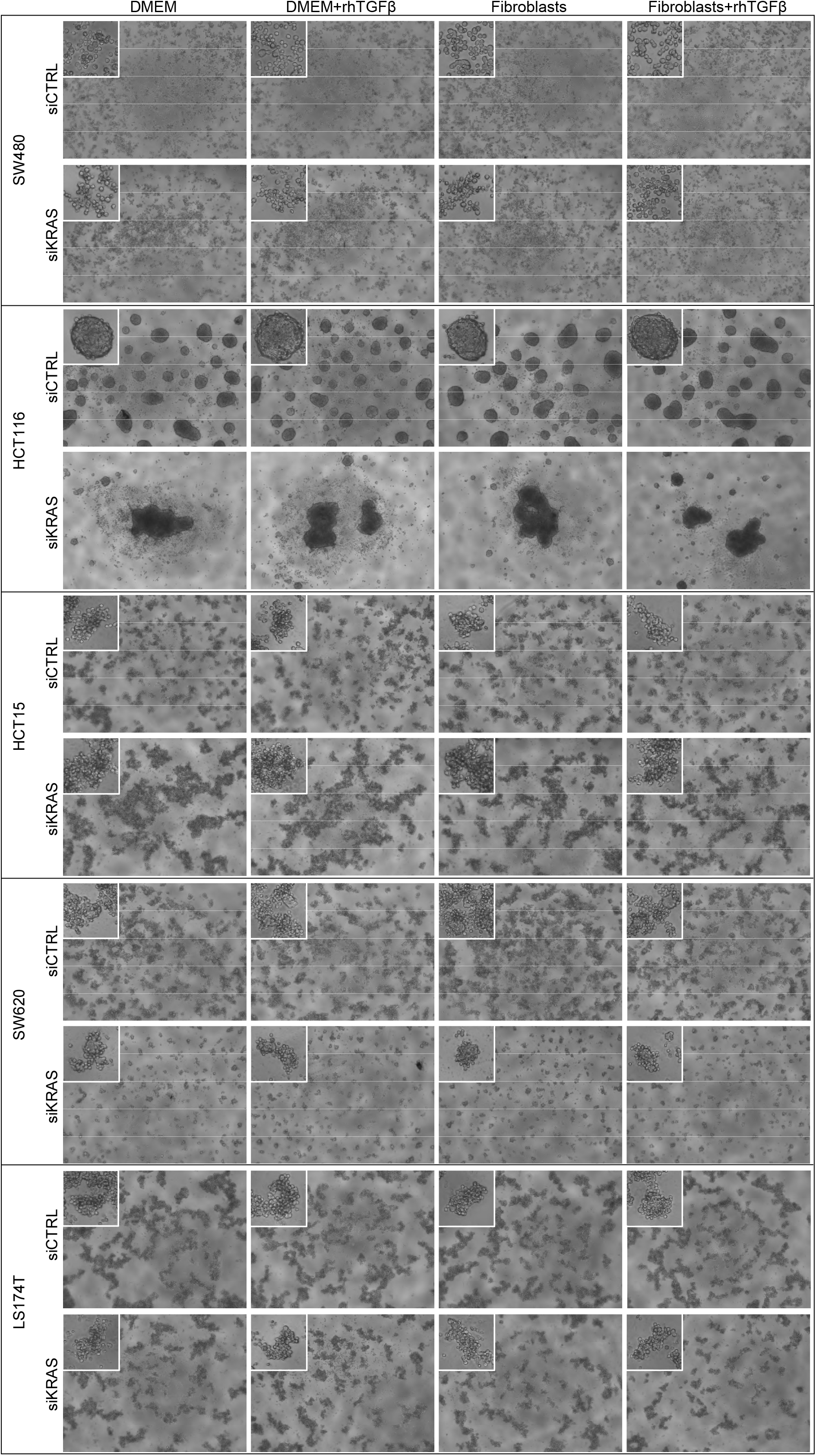
Fibroblast CM shows no impact on cell-cell aggregation capacity. Cell-cell aggregation capacity was evaluated in cells cultured in CM during 48 hours. SW480 cells, both siCTRL and siKRAS, lack the capacity to form aggregates. HCT116 and HCT15 siKRAS cells form larger aggregates when compared to siCTRL. On the other hand, SW620 siKRAS cells form smaller, yet more compact, aggregates when compared to siCTRL. LS174T do not show major differences between conditions. Representative images of 3 independent biological replicates, all obtained with a 5x objective, and a cropped aggregate (all of the same size) are shown for each condition.

To study the impact of fibroblasts-secreted factors on KRAS silenced cells invasion, siCTRL and siKRAS cells were subjected to a Matrigel invasion assay, using fibroblasts CM as a chemoattractant. Of note, we have observed that fibroblasts-induced invasion was independent of the state of fibroblasts activation. The results showed that the cell lines can be grouped according to their intrinsic/induced invasive capacity in response to the CM and to the effect of KRAS silencing (Fig. 3). For instance, SW480 cells invaded in all conditions independently of the CM. Its intrinsic capacity to invade depends on KRAS as siKRAS cells are less invasive. HCT116, HCT15 and SW620 cells increased invasion only when stimulated with fibroblasts CM. Importantly, in all these cell lines, siKRAS impaired fibroblast-induced invasion. LS174T cells have low basal invasion levels and did not become invasive when stimulated with CM. Surprisingly, in this cell line, siKRAS resulted in a significant increase of the invasive capacity in response to the fibroblasts CM.

**Figure 3.**
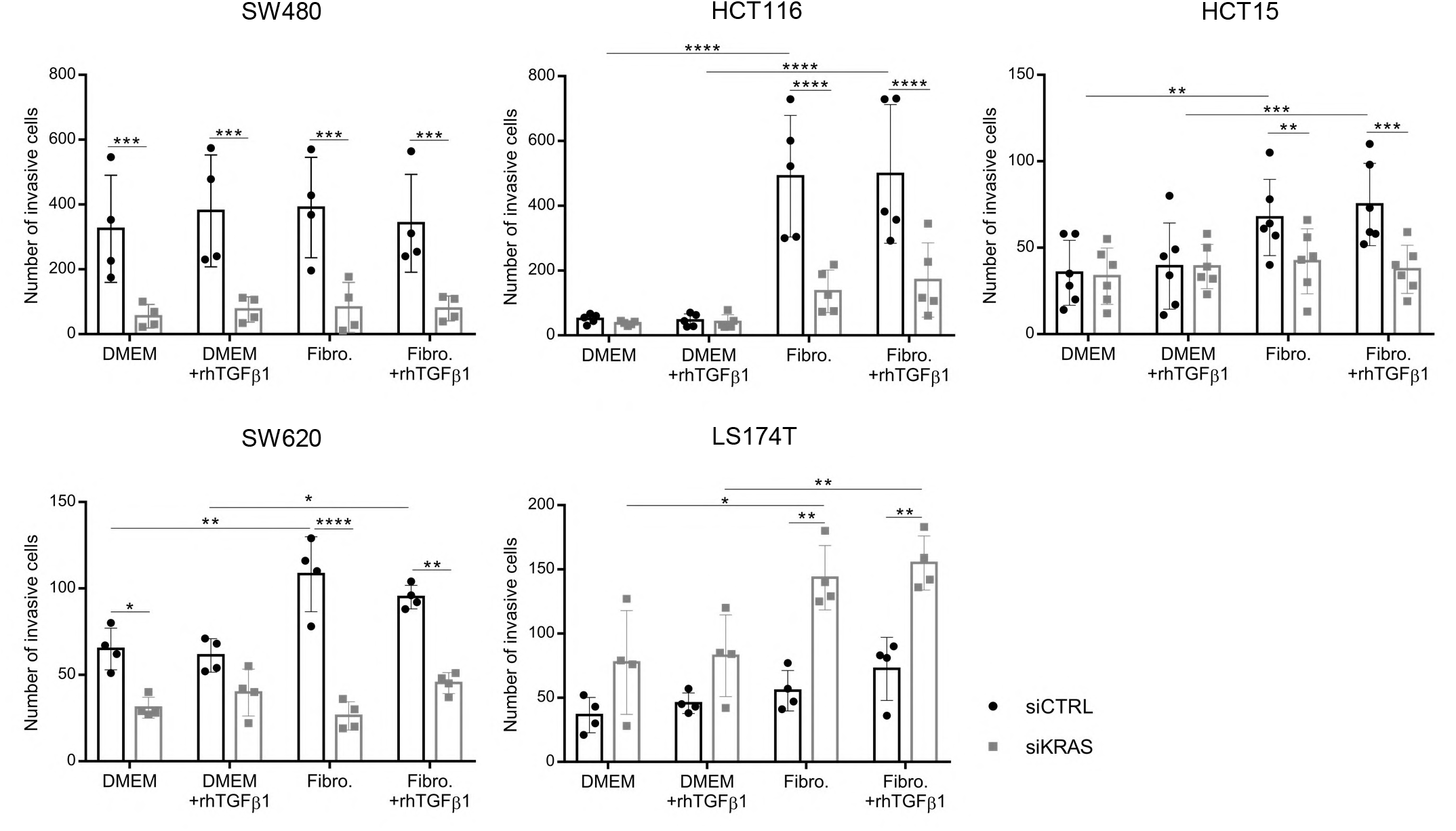
Mutant KRAS modulates fibroblast-induced colorectal cancer cell invasion in a cell line dependent manner. CM from normal-like (Fibro.) and rhTGFβ1-activated fibroblasts (Fibro+rhTGFβ1) and the respective controls (DMEM and DMEM+rhTGFβ1) were used as chemoattractant in Matrigel invasion assay. SW480 cells (siCTRL) invade independently of the CM and KRAS silencing (siKRAS) was capable to impair the invasion capacity. HCT116, HCT15 and SW620 cells (siCTRL) respond to the fibroblast CM increasing the invasion levels and KRAS silencing (siKRAS) impaired the invasive capacity. LS174T cells (siCTRL) displayed low levels of invasion and did not respond to the fibroblast CM. In this cell line, KRAS silencing (siKRAS) resulted in an increased invasion capacity in response to the fibroblast CM. Individual values for each biological replicate are plotted in the graph together with mean ±SD lines. Statistical significance was evaluated using the two-way ANOVA considering repeated measures by both factors, with Tukey’s multi comparison test (*p≤0.05; **p≤0.01; ***p≤0.001; ****p≤0.0001).

Together, these data showed that mutant KRAS exerts both fibroblasts secreted factors-independent and -dependent functions, depending on the cell line and on the effect. Specifically, cell cycle, and cell-cell aggregation of CRC cells were mainly regulated by KRAS through an autonomous signaling mechanism, therefore independent of fibroblasts-secreted factors. In the case of invasion, the results revealed that the type of response is cell line-dependent. Mutant KRAS can provide pro-invasion signals by itself but it can also function as a key modulator of cancer cell response to fibroblasts-derived pro-invasive factors. Noteworthy, we also showed that the inhibition of KRAS oncogene can promote cancer cell invasive capacity that is dependent on the stimulation with fibroblasts-derived factors.

### Fibroblast-induced invasion depends on the HGF-C-MET axis

From the previously analyzed cellular effects, invasion was the one that revealed more dependency on the fibroblasts-secreted factors. To further explore the molecular mechanism underlying the role of mutant KRAS in mediating fibroblasts-induced invasion, we selected the cell lines HCT116 in which KRAS silencing decreased fibroblasts-mediated invasion, and LS174T in which KRAS silencing increased fibroblasts-mediated invasion. For these experiments, we only used CM from rhTGFβ1-activated fibroblasts (and the respective control), as similar invasion levels were observed with both CM from non-activated and activated-fibroblasts.

Starting to explore the molecular mechanism underlying inhibition of invasion upon KRAS silencing, we performed a phospho-receptor tyrosine kinase (RTK) array aiming to obtain a more comprehensive view of the receptors involved in the process and to infer on key fibroblasts-secreted factors that could be mediating the effect. The results revealed an increase in the phosphorylation of the mesenchymal–epithelial transition factor (C-MET) in the invasive condition (HCT116 siCTRL treated with Fibro.+rhTGFβ1 CM) compared to the non-invasive ones (Fig 4A). Evaluation of the levels of the C-MET ligand, the hepatocyte growth factor (HGF), in the CM showed that non-activated and activated-fibroblasts secrete similar levels of this growth factor (Fig. 4B), a finding that is in accordance with the similar invasion levels observed in response to both media. In line with these observations, others have also detected similar HGF production in human colon derived fibroblasts and myofibroblasts (24). To verify whether HGF is indeed the driver of fibroblast-induced invasion, we evaluated the invasion levels in response to control medium supplemented with rhHGF, and HGF neutralization in the activated-fibroblasts CM. HGF supplementation resulted in an increased invasion capacity whereas HGF neutralization mimicked KRAS silencing, thus reducing the invasion capacity of HCT116 cells to the levels observed in the control medium (Fig. 4C). In addition, the analysis of C-MET expression showed that KRAS-silencing resulted in a downregulation of this receptor in all the analyzed cell lines, except for SW480 cells that expressed residual levels of C-MET (Fig. 4D and Supplementary Fig. S4). C-MET silencing resulted in a decreased invasive capacity of HCT116 cells in response to the CM, further proving the relevance of C-MET signaling in the invasive response (Fig 4E). This observation was also validated in SW620 cells (Supplementary Fig. S5). Together, these results demonstrate that fibroblast-induced invasion depend on the HGF-C-MET axis. Moreover, according to our knowledge, they award KRAS a role in mediating this process through the regulation of C-MET expression, for the first time.

**Figure 4.**
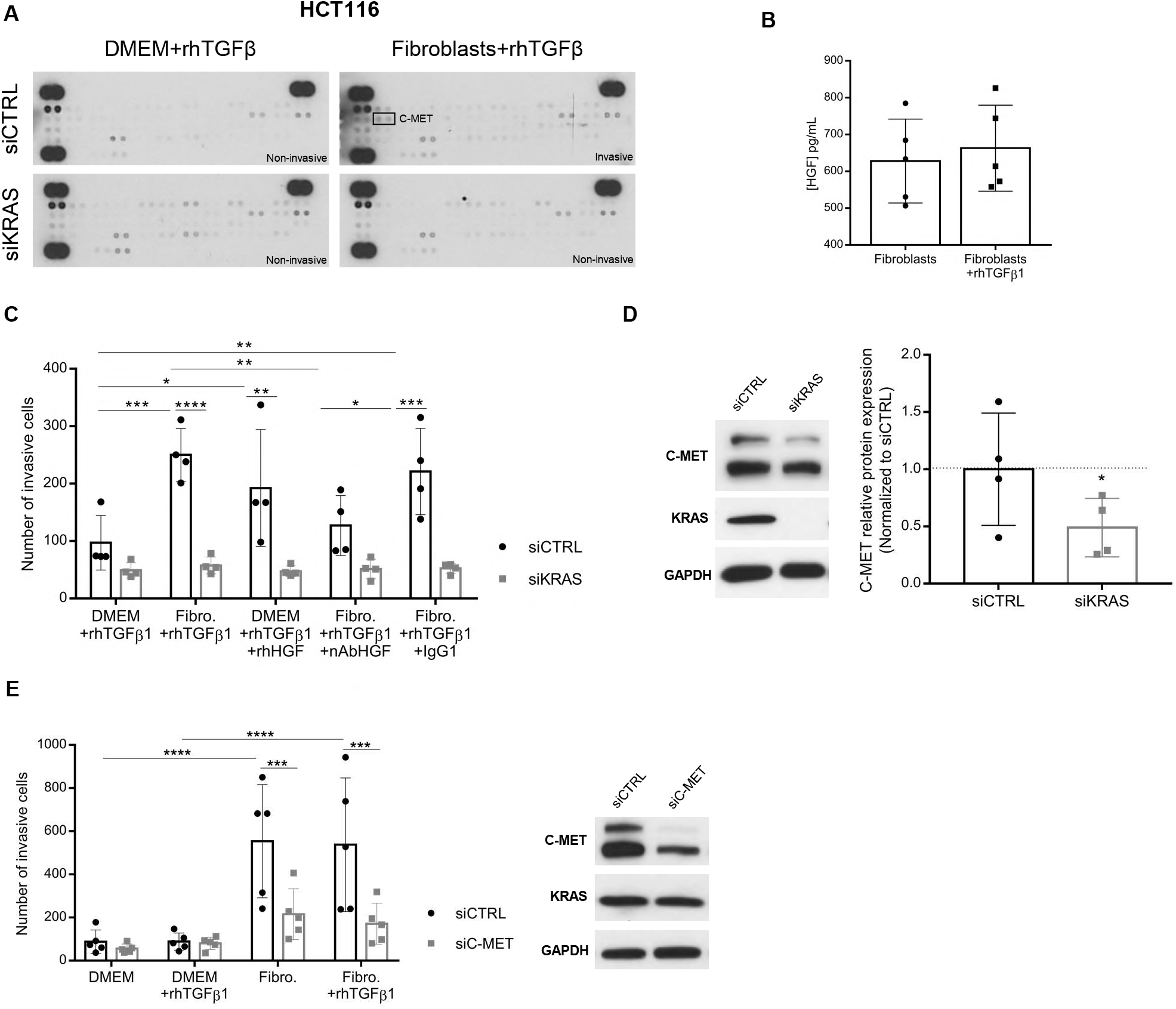
In HCT116 cells, fibroblast-induced invasion depends on the HGF-C-MET axis. **A,** HCT116 siCTRL and siKRAS cells were treated with Fibro.+rhTGFβ1 conditioned media and the respective control (DMEM+rhTGFβ1) for 24 hours and a Phospho-RTK array was performed. When comparing non-invasive with the invasive condition, an increase in C-MET phosphorylation is highlighted. **B,** Secreted HGF was quantified, by ELISA, in the conditioned media. Fibroblasts conditioned media showed to be enriched in HGF independently of the state of activation. **C,** Invasion of HCT116 cells (siCTRL and siKRAS) was evaluated upon stimulation with 50ng/mL of rhHGF (DMEM+rhTGFβ1+rhHGF), and HGF neutralization in the activated-fibroblasts conditioned medium (Fibro.+rhTGFβ1+nAbHGF). Fibro.+rhTGFβ1+IgG1 was used as control for the neutralizing antibody treatment. rhHGF stimulation resulted in an increased invasion capacity and HGF neutralization reduces the invasion capacity to similar levels observed in the control medium. KRAS silencing impaired the invasive capacity. **D,** KRAS silencing resulted in a downregulation of C-MET protein expression. **E,** Similarly to KRAS, C-MET silencing resulted in an impairment of the invasion capacity of HCT116 cells (siCTRL) in response to the fibroblast CM, in comparison to control CM conditions. Individual values for each biological replicate are plotted in the graph together with mean ±SD lines. Statistical significance was evaluated using the paired t-test (B,D) or the two-way ANOVA (C,E) considering repeated measures by both factors, with Tukey’s multi comparison test (*p≤0.05; **p≤0.01; ***p≤0.001; ****p≤0.0001).

Aiming to explore the mechanism behind the unexpected increase on the invasive capacity of LS174T cells upon KRAS-silencing, we first questioned whether this effect was specific of silencing KRAS or whether the inhibition of a similar oncogene would also trigger invasion. Since LS174T cells harbor an activating mutation at PIK3CA gene (25), we analyzed the invasive capacity of PIK3CA inhibited cells in response to CM from rhTGFβ1-activated fibroblasts. PIK3CA silencing had no impact on the invasive response either stimulated or not with fibroblast CM, demonstrating that the effect is specific of KRAS-silencing. Moreover, simultaneous inhibition of KRAS and PIK3CA was not enough to abolish invasion induced by KRAS-silencing (Fig. 5A), indicating that the presence of an hyperactivated form of PIK3CA does not function as an alternative pro-invasion pathway in the absence of KRAS. Noteworthy, despite no effect on the invasion capacity, the simultaneous silencing of KRAS and PIK3CA significantly decreased the number of cells, unlike their individual silencing where the number was unaltered (Supplementary Fig. S6).

**Figure 5.**
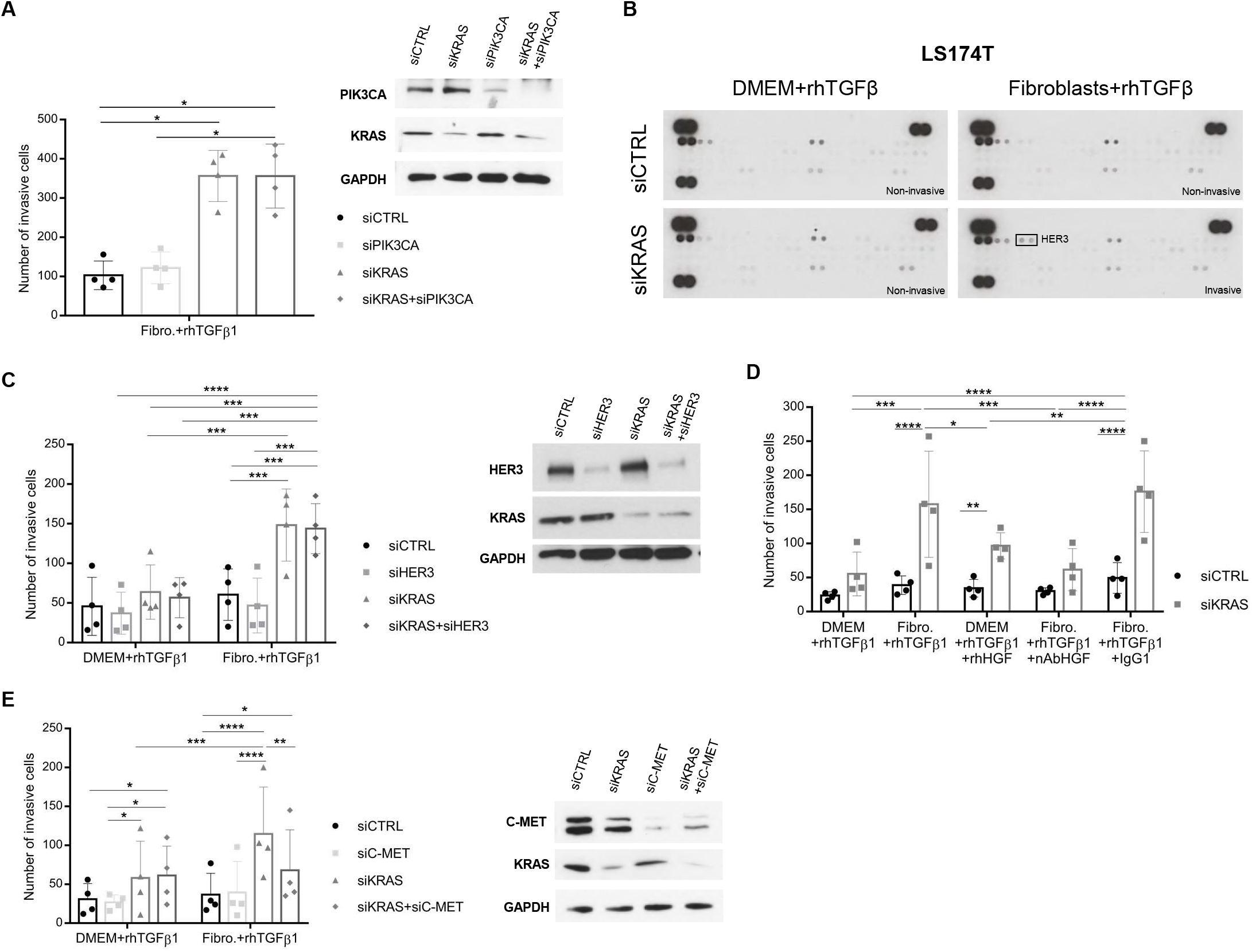
KRAS-silencing induced invasion of LS174T cells is impaired upon KRAS and C-MET simultaneous silencing. **A,**. Using Fibro.+rhTGFβ1 CM as chemoattractant, LS174T cells with KRAS and/or PIK3CA silenced were subjected to Matrigel invasion assay. PIK3CA silencing alone, either stimulated or not with Fibro.+rhTGFβ1 CM, had no impact on the invasive capacity, when compared to siCTRL cells. Simultaneous silencing of KRAS and PIK3CA was not enough to abrogate the increased invasion levels caused by KRAS silencing. Representative western blot showing silencing efficiency **B,** LS174T siCTRL and siKRAS cells were treated with Fibro.+rhTGFβ1 CM and the respective control (DMEM+rhTGFβ1) for 24 hours and a Phospho-RTK array was performed. When comparing non-invasive with the invasive condition, an increase in HER3 phosphorylation is highlighted. **C,** Using Fibro.+rhTGFβ1 CM and the respective control (DMEM+rhTGFβ1) as chemoattractant, LS174T with KRAS and/or HER3 silenced were subjected to Matrigel invasion assay. HER3 silencing alone had no impact on the invasive capacity of LS174T, when compared to siCTRL cells. Simultaneous silencing of KRAS and HER3 was not enough to abrogate the increased invasion levels caused by KRAS silencing. Representative western blot showing silencing efficiency. **D,** Invasion of LS174T siCTRL and siKRAS cells was evaluated upon stimulation with 50ng/mL of HGF (DMEM+rhTGFβ1+rhHGF), and HGF neutralization in the activated-fibroblasts conditioned medium (Fibro.+rhTGFβ1+nAbHGF). Fibro.+rhTGFβ1+IgG1 was used as control for the neutralizing antibody treatment. Upon KRAS silencing, rhHGF stimulation resulted in an increase of the invasion capacity that is still significatively different than the levels induced by Fibro.+rhTGFβ1 CM. HGF neutralization reduces the invasion capacity. **E,** Using Fibro.+rhTGFβ1 CM and the respective control (DMEM+rhTGFβ1) as chemoattractant, LS174T with KRAS and/or C-MET silenced were subjected to Matrigel invasion assay. C-MET silencing alone demonstrated no impact on the invasive capacity of LS174T, when compared to siCTRL cells. Simultaneous silencing of KRAS and C-MET decreased the levels of KRAS-silencing induced invasion. Representative western blot showing silencing efficiency. Individual values for each biological replicate are plotted in the graph together with mean ±SD lines. Statistical significance was evaluated using the one-way (A) two-way ANOVA (C,D,E) considering repeated measures by both factors, with Tukey’s multi comparison test (*p≤0.05; **p≤0.01; ***p≤0.001; ****p≤0.0001).

Since LS174T cells are described to belong to the metabolic deregulated consensus molecular subtype 3 (26), we postulated that siKRAS could induce a metabolic imbalance, prompting cells to move towards external sources of energy. Considering that fibroblast-induced invasion of CRC cells can be driven by growth factor-independent factors such as lipids, we followed a published protocol to deplete proteins from the rhTGFβ1-activated fibroblasts CM while keeping small metabolites (23). The results showed that protein-depleted CM was not capable of inducing invasion of KRAS-silenced cells, implying that invasion of LS174T siKRAS cells is driven by a protein-like factor (Supplementary Fig. S7). Therefore, we performed a phospho-RTK array which revealed an increase in HER3 phosphorylation on the invasive condition (KRAS-silenced cells stimulated with Fibro.+rhTGFβ1 CM) (Fig. 5B). To validate HER3 as a mediator of siKRAS invasive response, we silenced its expression using siRNA. HER3 silencing alone did not induce any alterations in the invasive capacity of LS174T cells, either stimulated or not with fibroblast CM. Likewise, simultaneous silencing of KRAS and HER3 did not abrogate the increased invasion capacity induced by KRAS-silencing (Fig. 5C). Therefore, we excluded HER3 as the main mediator of LS174T invasive response. Considering that KRAS silencing-induced invasion occurred in response to CM from both non-activated and activated fibroblasts, and that both CM displayed similar levels of HGF, we decided to explore the role of HGF in this context. Chemoattraction with control CM supplemented with rhHGF increased the number of siKRAS invasive cells that, despite being significatively higher than control cells, was still smaller than the levels observed in response to Fibro+rhTGFβ1 CM. Moreover, HGF neutralization reduced the invasion capacity of siKRAS cells to similar levels observed in the control medium (Fig. 5D). Since we observed a partial response to HGF stimulation, we sought C-MET inhibition. The results showed that C-MET silencing alone had no effect on the invasive capacity, while in combination with KRAS silencing, it reduced invasion (Fig 5E). Altogether, the results on LS174T cells suggest that the increased invasive response, seems to be a KRAS-silencing specific effect, also mediated by the HGF/C-MET axis.

Overall, these data show that despite the cell line-specific invasion responses of siKRAS cells, the HGF-C-MET axis seem to be a common player.

## Discussion

Our work highlights that KRAS simultaneously exerts autonomous and non-autonomous effects in CRC cells, further reinforcing the magnitude and complexity of oncogenic KRAS signaling. On the one hand, we showed that cell cycle progression and cell-cell aggregation are mainly controlled by KRAS, independently of external stimulation with fibroblasts-derived factors. On the other hand, the work awards oncogenic KRAS an important role on the modulation of the response to fibroblast-secreted factors, contributing to understand the mechanisms driving CRC cells invasion.

Herein we also evidenced that CRC cell lines, that poorly invade at basal levels, become invasive in response to fibroblast-secreted HGF. Mechanistically, we show that, in mutant KRAS CRC cell lines that invade in response to fibroblasts CM, KRAS silencing impairs HGF-driven invasion by downregulating C-MET expression (Fig.6A). The observed link between KRAS, HGF/C-MET axis and invasion of CRC cells is supported by the literature. HGF has long been recognized as a fibroblast-derived pro-invasive factor (24,27). In addition, early *in vitro* and *in vivo* studies addressing the role of HGF-C-MET signaling in the context of RAS oncogenic activation have uncovered that HRAS transformation resulted in consequent C-MET overexpression (28,29) which played an essential role in tumor growth and metastasis (30). In a KRAS mutant CRC model, C-MET overexpression-mediated tumorigenesis was found to be dependent on oncogenic KRAS activation and not on its wild-type counterpart (31). In human CRC samples, C-MET overexpression is well documented, being a predictor of tumor invasion and regional metastasis, and a biomarker of poor prognosis (32–34). Likewise, HGF levels in cancer tissue and in serum have also been shown to be elevated in CRC patients. Serum levels correlated with tumor size, lymph-node involvement, and distant metastasis, being an independent risk factor for poor survival in Stage II/III. Moreover, C-Met and HGF co-expression is correlated with advanced stage and poor survival, predicting a metastatic phenotype (35). Additionally, the evaluation of C-MET levels in surgical resections of untreated KRAS mutant and KRAS wild-type CRC cases revealed that KRAS mutant tumors have higher C-MET score (36). Together with ours, these works show the relevance of the HGF-C-MET axis in RAS-driven tumorigenesis. Importantly, representative studies evaluating the relevance of C-MET and HGF expression in the context of mutant KRAS CRC are needed to properly assess the potential prognostic and biomarker applications in this subset of cases.

To the best of our knowledge, we report for the first time, that KRAS silencing may also provide pro-invasive signals. This unexpected result found in LS174T cell line, seems to be specific from KRAS silencing as the inhibition of PIK3CA oncogene does not promote invasion. Noteworthy, despite failing to show an effect in the invasive capacity, simultaneous silencing of KRAS and PIK3CA resulted in a significant reduction in the number of cells. In this context, although mutant KRAS-inhibited cells would invade more, the simultaneous inhibition of both oncogenes would blunt tumor outgrowth from KRAS-silenced invasive cells, suggesting a promising anti-tumorigenic effect for a combined therapy. In addition to these observations, we also attempted to dissect the molecular mechanism underlying the gain of invasive capacity upon KRAS silencing as this could hint on possible mechanisms of resistance to KRAS inhibitors and reveal potential combinatorial therapeutic opportunities. Through different approaches, we demonstrated that although KRAS silencing decreases C-MET expression, the pro-invasive signals seem to partially depend on fibroblasts-derived HGF and on the remaining C-MET. Whether other RTK cooperate with C-MET and whether C-MET acts as a homodimer or if it takes advantage of its capacity to heterodimerize with different cell surface proteins (37) demands further clarification. The plasticity of C-MET heterodimerization with other RTK, for instance with another member of the HER family, may explain why HER3 inhibition did not induce any effect on invasive capacity of KRAS-inhibited cells (Fig. 6B). Still, why LS174T cells invade in response to fibroblasts-derived HGF only upon KRAS inhibition remains an open question. Data generated from recent studies addressing the resistance mechanisms to KRAS G12C specific inhibitor has revealed that KRAS inhibition releases cells from the negative feedback loops that normally restrain RTK and MAPK activation, triggering a rapid activation of multiple RTK and downstream signaling – vertical pathway activation (38). Therefore, we speculate that a similar mechanism may be occurring in our model: mutant KRAS signaling maintains active the feedback inhibitory loop that blocks signaling from fibroblasts through restraining the activation of pathways downstream to C-MET and other RTKs; when KRAS is silenced, the negative feedback regulation is lost, allowing the activation of cell surface receptors and downstream signaling pathways that drive invasion (Fig. 6B). Furthermore, despite harboring a KRAS mutation, the LS174T cell line is classified as KRAS independent (39), meaning that these cells do not depend on KRAS signaling to maintain their malignant features.

**Figure 6.**
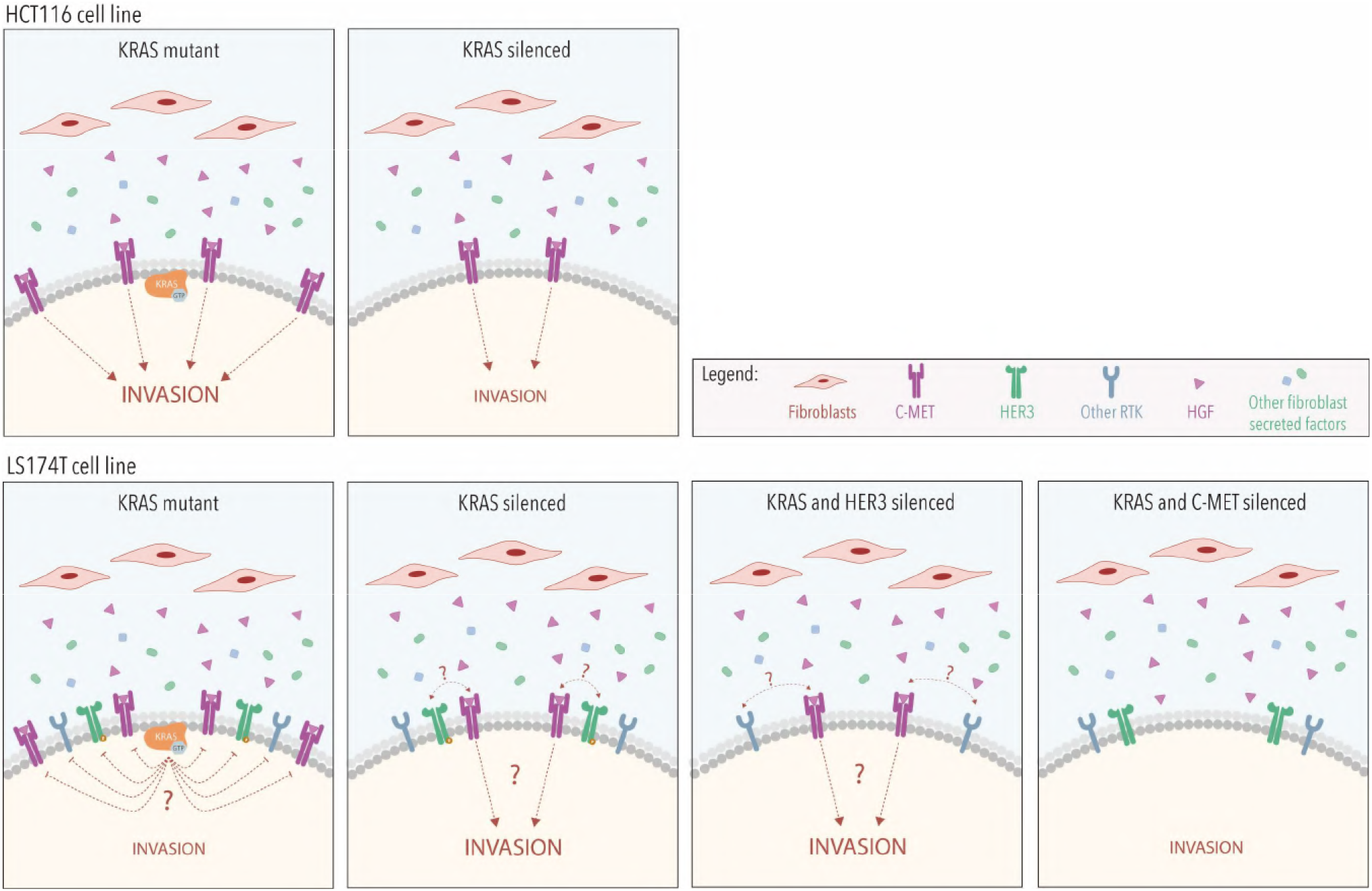
Proposed model underlying the invasive responses. In HCT116 cell line, fibroblast-secreted HGF binds C-MET driving the invasive response. When KRAS is silenced, C-MET expression is downregulated resulting in reduced invasion levels. In LS174T, the cell line that invades in response to fibroblast CM only when KRAS is silenced, we hypothesize that mutant KRAS signaling activates the negative feedback loops to restrain the level of receptor tyrosine kinase (RTKs) activation. Therefore, upon KRAS loss, a number of RTKs are released from the negative feedback becoming free to activate other signaling pathways, driving the invasive response. Owing to the lack of effect of HER3 inhibition and to the partial response to HGF, we speculate that C-MET is heterodimerizing with other RTK to induce the invasive response. Simultaneous silencing of KRAS and C-MET results in the decrease of the invasive capacity, thus proving the key role of C-MET in this process.

Therefore, KRAS loss is not detrimental to these cells but by releasing the brakes from RTK, it would facilitate invasion through KRAS-independent signaling pathways. Coincidently, in the study where the vertical pathway activation effect is described, the two mutant KRAS CRC cell lines used are also considered mutant KRAS-independent (38). Therefore, other KRAS independent cell lines, including the ones harboring KRAS G12C mutations, should be evaluated to assess if this classification justifies the response that we observed. Moreover, the discovery of biomarkers that could discriminate the two group of responses would have great value to stratify the patients that would indeed benefit from a KRAS-targeting therapy.

Notwithstanding the opposite effects observed upon KRAS silencing, the HGF-C-MET axis demonstrated to be a main player in both scenarios, serving as a rationale to explore this axis as a possible therapeutic target in the context of KRAS mutant CRC, with particular emphasis in combinatorial treatment regimens. Of note, in *in vitro* and *in vivo* models, stroma-derived HGF was found to underly the resistance to anti-angiogenic agents by promoting the metabolic adaptation of CRC cells (40). Since the antiangiogenic agents figure on the first and second-line treatment options in cases of KRAS mutant CRC (2), the combination with a C-MET/HGF inhibitor seems rational. Clinically, targeting the KRAS-C-MET-HGF axis can be particularly relevant in the setting of CRC liver metastasis (CLM). In this scenario, the presence of KRAS mutations is associated with poor survival, shorter disease-free interval from primary tumor resection to detection of liver metastasis and presence of multiple metastasis (41). Additionally, in patients who underwent curative resection of CLM, besides being associated with worse overall survival and recurrence-free survival, (K or N) RAS mutation predicts early lung recurrence after curative resection (42). Mutant RAS cells exhibit an infiltrative growth pattern and the mutant status associates with wider positive surgical margins compared to RAS wild-type tumors (43). Therefore, it has been suggested that resection of mutant RAS CLM should be approached cautiously (44,45). Interestingly, the liver is not the preferential site for mutant RAS CRC cells to metastasize. Nevertheless, RAS mutations are more frequently found in lung and peritoneal metastases than in the liver. Yet, the poor prognostic impact of mutant RAS seems to be specific for patients undergoing resection of liver metastases as it did not affect survival after removal of lung or peritoneal metastases (46,47). The reason why this happens is yet to be identified. However, our data linking fibroblasts-derived HGF/C-MET and mutant KRAS pinpoints a possible molecular mechanism underlying the adverse impact of KRAS mutations in the prognosis of patients undergoing CLM resection. Moreover, in an *in vivo* model, CLM resection was found to be followed by C-MET upregulation, as a result of the liver regeneration process, and consequent tumor growth (48). HGF plasma levels have also found significatively elevated after hepatectomy (49), in accordance with its function as an essential factor for liver regeneration. Given all these evidence pointing to a tumor promoting environment in the liver upon surgery, the validation of the KRAS-C-MET-HGF axis in this setting can help to stratify the patients and better treat the ones that could benefit from an adjuvant anti-C-MET/HGF therapy.

In summary, this work highlights the many faces of KRAS oncogenic signaling and contributes to the understanding of the mechanisms underlying the response of KRAS mutant cancer cells to microenvironmental cues. Furthermore, it reinforces the idea that microenvironment-derived signals should be taken into consideration for combinatorial treatments, as they are key players in the resistance mechanism to KRAS inhibition. Importantly, we disclosed the HGF-C-MET axis a possible therapeutic target deserving further attention in the context of KRAS mutant CRC cases, which currently lack efficient therapies.

## Notes

The authors declare no potential conflict of interests.

### Competing Interest Statement

The authors have declared no competing interest.

