## Supplementary files for "Mutant KRAS modulates colorectal cancer cells invasive response to fibroblast-secreted factors through the HGF/C-MET axis"

### Supplementary Figures and Tables

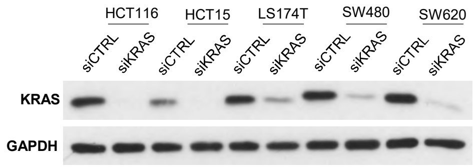


**Figure S1. Representative western blot illustrating KRAS silencing efficiency in all cell lines.**

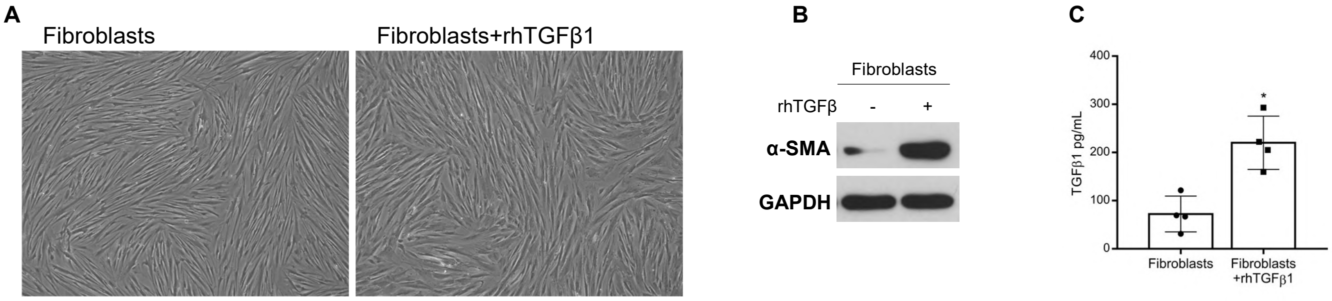


**Figure S2. rhTGFβ1 successfully activates CCD18-Co colon fibroblasts**. **A.** Representative micrographs (5x objective) showing phenotypical differences between Fibroblasts and Fibroblasts activated with rhTGFβ1. **B.** Representative western blot showing increased a-SMA expression by rhTGFβ1-activated fibroblasts. **C.** Activated fibroblasts secrete significatively more TGFβ1 than “normal-like” fibroblasts.

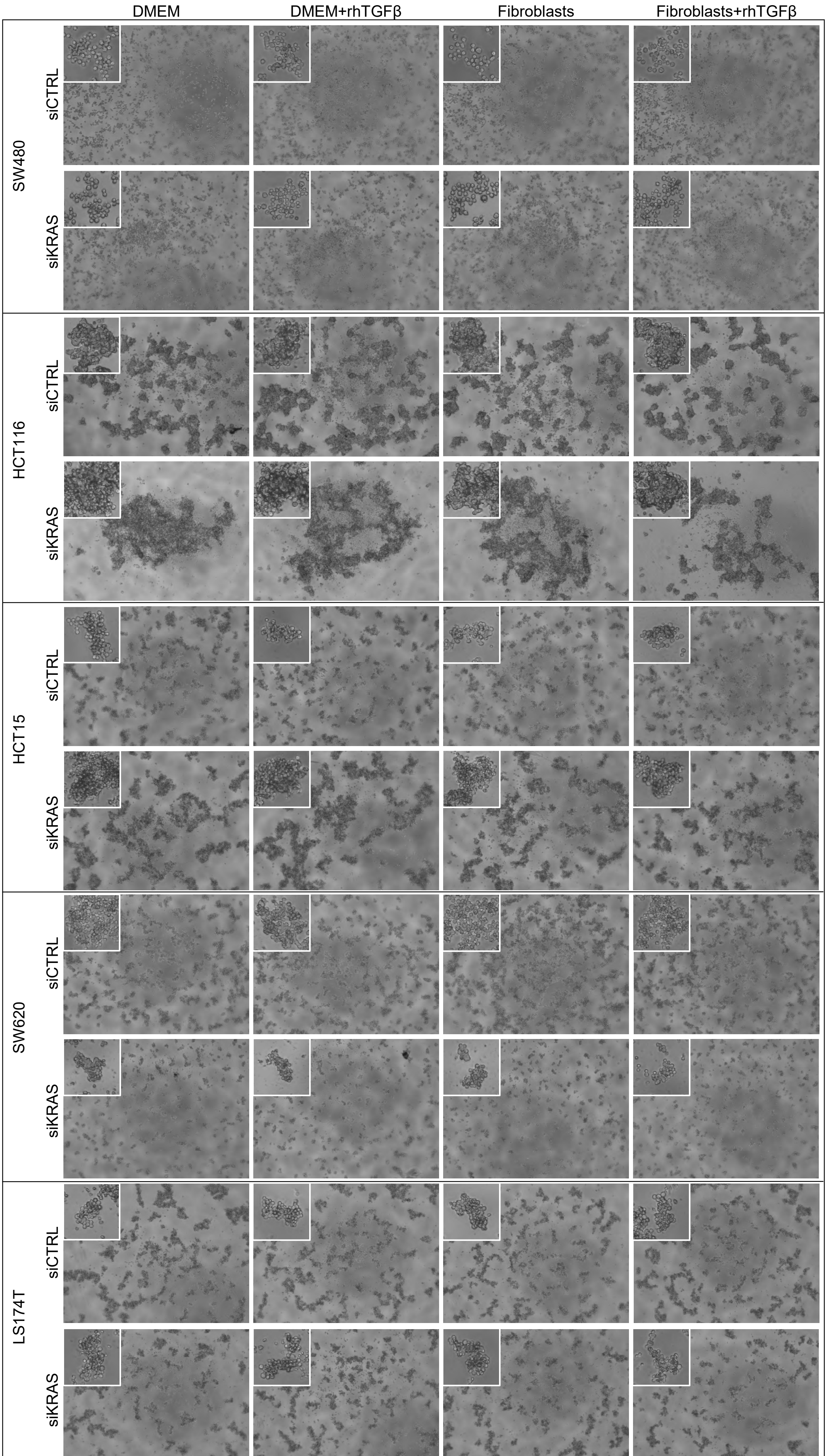
DMEM
DMEM+rhTGFβ
Fibroblasts
Fibroblasts+rhTGFβ
SW480
siCTRL
siKRAS
HCT116
siCTRL
siKRAS
HCT15
siCTRL
siKRAS
SW620
siCTRL
siKRAS
LS174T
siCTRL
siKRAS

**Figure S3. Fibroblast CM shows no impact on cell-cell aggregation capacity.** Cell-cell aggregation capacity was evaluated in cells cultured in CM during 24 hours. SW480 cells, both siCTRL and siKRAS, lack the capacity to form aggregates. HCT116 and HCT15 siKRAS cells form larger aggregates when compared to siCTRL. On the other hand, SW620 siKRAS cells form smaller, yet more compact, aggregates when compared to siCTRL. LS174T do not show major differences between conditions. Representative images of 3 independent biological replicates, all obtained with the 5x objective, and a cropped aggregate (all of the same size) are shown for each condition.

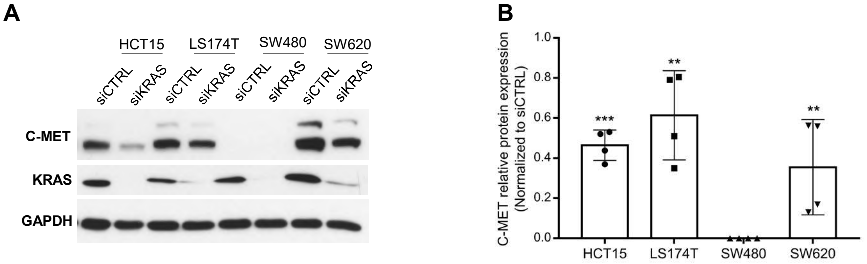


**Figure S4. KRAS silencing results in the downregulation of C-MET expression. A.** Representative western blot showing decreased C-MET expression upon KRAS inhibition in all cell lines, with the exception of SW480 that express residual levels of this receptor. **B.** Western blot quantification of four independent experiments showing significant C-MET downregulation. Values were normalized and compared to siCTRL condition using *t-test* (**p≤0.01; ***p≤0.001).

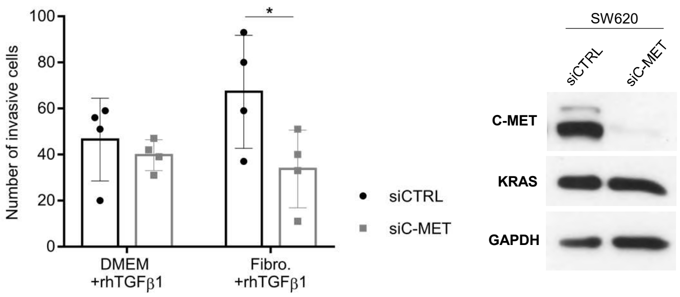


**Figure S5. C-MET silencing decreases the invasive capacity of SW620 cells.** Statistical significance was evaluated using the two-way ANOVA considering repeated measures by both factors, with Tukey's multi comparison test (*p≤0.05).

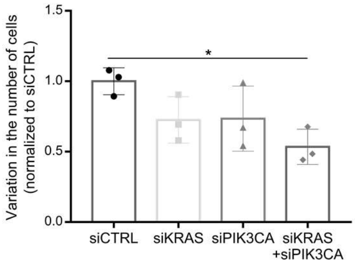


**Figure S6. Simultaneous silencing of KRAS and PIK3CA in LS174T cells results in a reduction in the number of cells.** Statistical significance was evaluated using the one-way ANOVA, with Tukey's multi comparison test (*p≤0.05).

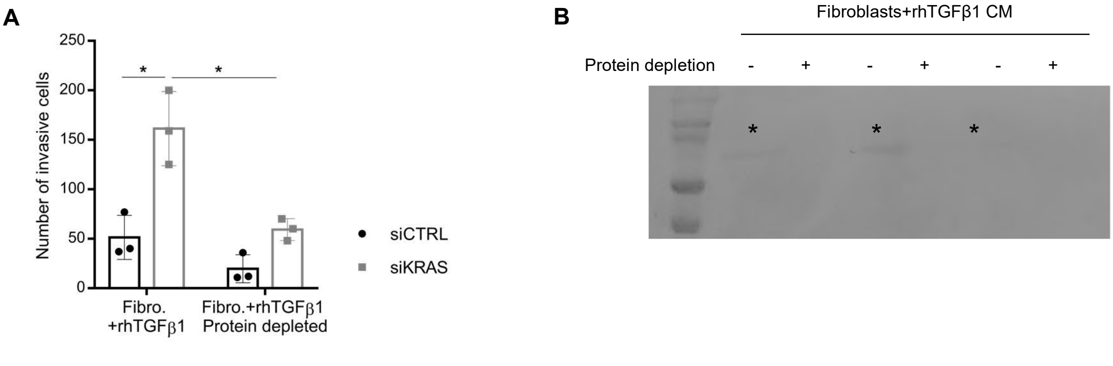


**Figure S7. In LS174T cells, KRAS silencing induced invasion is dependent on the presence of protein factors. A,** Normal and protein depleted Fibroblasts+rhTGFβ1 conditioned medium were used as chemoattractant in Matrigel invasion assay using LS174T siCTRL and siKRAS cells. KRAS-silencing induced invasion was significantly decreased upon CM protein depletion. Statistical significance was evaluated using the two-way ANOVA considering repeated measures by both factors, with Tukey's multi comparison test (*p≤0.05). **B,** Paired Fibroblasts+rhTGFβ1 CM subjected or not to protein depletion protocol were ran in an SDS-PAGE gel and transferred to a membrane that was stained with Ponceau S solution. The presence of visible bands (*) was only observed in non-depleted medium, being indicative of efficient protein-removal upon protein-depletion protocol.

**Supplementary Table S1.** Number of seeded cells for each cell line

| Cell line | Number of seeded cells/well (6-well plate) | |
|---|---|---|
| | For cell cycle analysis | Other assays |
| **HCT116** | 1×$10^5$ | 1.5×$10^5$ |
| **HCT15** | 1.5×$10^5$ | 2×$10^5$ |
| **SW480** | 1×$10^5$ | 1.5×$10^5$ |
| **SW620** | 3.5×$10^5$ | 4×$10^5$ |
| **LS174T** | 1,5×$10^5$ | 2×$10^5$ |

**Supplementary Table S2.** Primary antibodies used in western blot

| Antibody | Reference | Manufacturer | Dilution | Blocking agent |
|---|---|---|---|---|
| **KRAS** | LS-C175665 | LS-Bio | 1:4000 | 5% non-fat milk in PBS+0.5% Tween20 |
| **GAPDH** | sc-47724 | Santa Cruz Biothecnology | 1:10000 | 5% non-fat milk in PBS+0.5% Tween20 |
| **C-MET** | sc-10 | Santa Cruz Biothecnology | 1:1000 | 5% non-fat milk in PBS+0.5% Tween20 |
| **HER3** | 12708 | Cell signaling | 1:1000 | 5% non-fat milk in PBS+0.5% Tween20 |
| **PIK3CA** | 4249 | Cell signaling | 1:1000 | 4% BSA in PBS+0,5% Tween20 |
| **α-SMA** | ab7817 | Abcam | 1:250 | 5% non-fat milk in PBS+0.5% Tween20 |

**Supplementary Table S3.** CRC Cell lines used in this work and their main characteristics (MSI- Microsatellite instable; MSS- Microsatellite stable) (Adapted from Ahmed et al., 2013- reference 25)

| Cell line | MSI status | KRAS | PI3KCA | Derived from |
|---|---|---|---|---|
| **HCT116** | MSI | G13D | H1047R | Primary tumor |
| **HCT15** | MSI | G13D | E545K;D549N | HCT-15/DLD-1 misclassified |
| **SW480** | MSS | G12V | wt | Primary tumor |
| **SW620** | MSS | G12V | wt | Lymph node metastasis |
| **LS174T** | MSI | G12D | H1047R | Subcultured LS 180 |
